# Distinct representation of cognitive flexibility and habitual stability in the primate putamen, caudate, and ventral striatum

**DOI:** 10.1101/2024.02.27.579257

**Authors:** Shin-young An, Seong-Hwan Hwang, Keonwoo Lee, Hyoung F. Kim

## Abstract

Recent primate studies have demonstrated a functional distinction along the rostral—caudal axis of the striatum, which has challenged the conventional view that flexible adaptation and habitual action differ in processing along the medial—lateral axis. We found that neurons in the rostral putamen, caudate, and ventral striatum encode values flexibly updated for adaptive choices, rather than values stably sustained for visual habit. In the reversal value learning, rostral striatal neurons dynamically updated their value discrimination responses after value reversals, whereas, in the stable value retrieval, most did not encode the value. Notably, caudate neurons were faster to update values after reversal trials than ventral striatum neurons. Slow-learning neurons were identified selectively in the ventral striatum. In each trial, their learning speeds were similar during initial learning, suggesting a parallel value update in each striatal region. Our findings thus indicate that the rostral striatum prioritizes cognitive flexibility over habitual stability.

## Introduction

Cognitive flexibility is the ability to adapt to changes in task demands or rules by switching between mental sets, which allows animals to recognize the meaning of sensory stimuli even when they change, and then adjust their behavior to maximize their survival. This ability to switch between different mental sets is believed to rely on a specialized brain system responsible for recognizing the current meaning of stimuli and decision-making.

This cognitive flexibility can be evaluated by using the reversal task, a widely used experimental paradigm in animal and human research (*1*). The task typically consists of training an individual on a simple rule, such as associating a particular stimulus (e.g., a visual object) with a particular outcome (e.g., a liquid reward). Once the individual has learned the rule, it is then reversed, and the individual is required to switch to the new rule and respond accordingly. The ability to adapt to the rule change and respond correctly is considered an indicator of cognitive flexibility.

Past studies used this reversal task paradigm and found that certain brain regions in the cortical and subcortical areas, including the orbitofrontal cortex, medial prefrontal cortex, amygdala, and caudate nucleus, are involved in cognitive flexibility (*1–4*). Interestingly, it has been found that a specific area within a structure plays a selective role in cognitive flexibility. Past studies in rodent models demonstrated functional distinctions along the medial—lateral axis of the rodent striatum (*5*). The dorsomedial striatum (DMS) plays a role in flexible goal-directed behavior, whereas the dorsolateral striatum (DLS) is primarily involved in habitual behavior. It is therefore widely accepted that these structures in rodents, the DMS, and DLS, are anatomically homologous to the caudate and putamen in primates, respectively (*5–9*). Several human fMRI and rodent studies have implicated the functional homologies of the caudate and DMS in goal-directed behavior and those of the putamen and DLS in the control of habitual actions (*10–12*).

However, findings in primate striatum research have raised a question about the validity of this idea (*13–15*). Notably, a marmoset study revealed that inactivation of the putamen impaired the reversal task, which suggests its role in cognitive flexibility (*13*). Furthermore, recent research in macaque monkeys revealed that goal-directed and habitual behavior are guided through distinct regions of the basal ganglia along the rostral-caudal axis, rather than the medial-lateral axis. For example, the rostral portion of the caudate (caudate head) is selectively involved in the reversal task, whereas the caudal part (caudate tail) plays a role in habitual behavior (*2*, *16–18*). These findings motivate the exploration of various primate striatal regions, considering their distinct characteristics compared to rodents, in the context of cognitive flexibility.

The anatomical regions that receive inputs from dopamine (DA) neurons provide insights into the striatal regions involved in cognitive flexibility because the reward prediction error (RPE) signal from these dopamine neurons is believed to play an important role in reversal learning (*19–22*). Notably, dopamine neurons signaling RPE in the rostral-medial SNc project not only to the caudate head but also to the putamen, suggesting its potential role in cognitive flexibility (*23–26*). In addition, the putamen receives inputs from the prefrontal cortex responsible for cognitive flexibility, as well as sensory and motor cortices involved in habitual behavior (*27*–*30*). While previous research has mainly focused on the role of caudate in cognitive flexibility, the neuronal encoding of cognitive flexibility and habitual stability in the primate putamen remains to be explored through direct neuronal recording.

The ventral striatum is another structure that receives dopaminergic inputs from the ventral tegmental area (VTA), encoding RPE signal (*31–34*). Recent studies have shown that neurons in the primate ventral striatum stably maintained previously learned object values for habitual behavior (*35*). However, their neural encoding of cognitive flexibility remains unclear in primates. Therefore, investigating whether and how individual neurons in these three striatal regions, the ventral striatum, caudate, and putamen, process flexibly changing and stably sustained values of visual objects is crucial for understanding the neural basis of cognitive flexibility and habitual stability.

In this study, we investigated the processing of cognitive flexibility and habitual stability by individual neurons in the primate caudate, putamen, and ventral striatum. Using single-unit recording, these individual neurons were examined while monkeys performed both a value-reversal task and a stable value-retrieval task. Our findings uncovered that individual neurons in the rostral striatum primarily encoded the dynamically changing values of visual objects, but did not exhibit encoding for stably sustained values. Furthermore, we analyzed the learning speed across these three striatal structures so as to explore how value information is processed across these structures.

## Results

### Flexible value-guided adaptive saccades and stable value-guided habitual saccades

To investigate the processing of different value information in the three striatal regions, we employed an object value-reversal task and a stable value-retrieval task following a long-term value learning procedure (Figure 1). In the object value-reversal task (Figure 1A), one of the two visual fractal objects was associated with a reward (high value), while the other was not (low value). This contingency was reversed between blocks. Their reaction times became significantly faster for high-valued objects compared to those for low-valued objects after a block switch (p < 0.001, two-tailed t-test) (Figures 1B and 4). As choice trials were introduced in the object value-reversal task, the monkeys demonstrated successful selection of a high-valued object after value reversals (Figures S1A and B). These observations indicated that the monkeys flexibly adapted their saccades in response to the reversed object-reward contingencies.

**Fig. 1.**
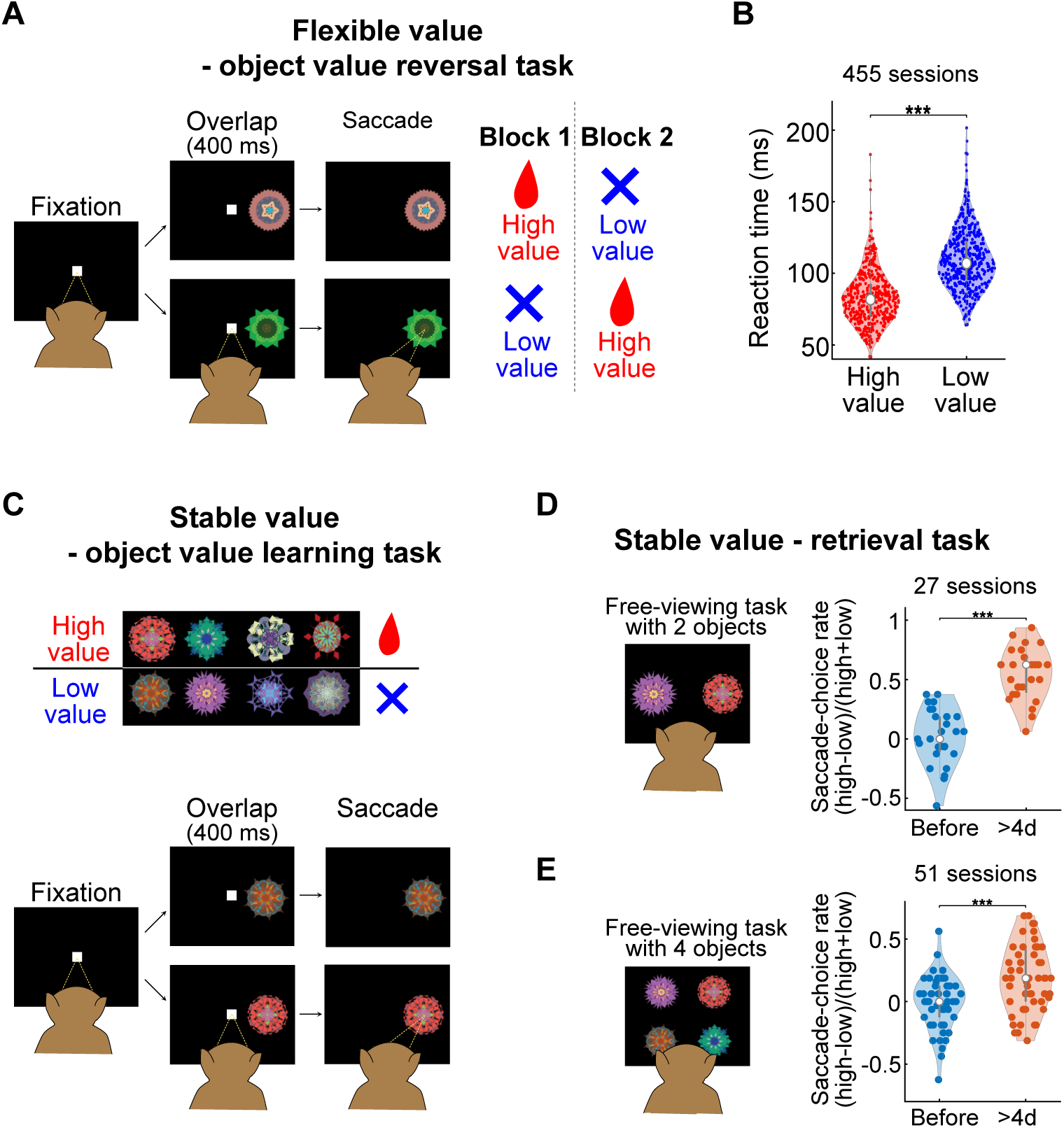
Behaviors guided by flexible and stable values. **(A)** The sequence of events in the object value-reversal task. While the monkey was fixating on a central white dot, one of the two objects appeared at the left or right position. After 400 ms, the fixation dot was turned off and the monkey was required to make a saccade to the object. In a block of 24–30 trials, the monkey received a liquid reward after making a saccade to one object, but not the other one. The object-reward association reversed in the subsequent block. **(B)** The average reaction times for high-valued and low-valued objects. The central white dot in a violin plot indicates the median, while the bottom and the top of the middle gray box correspond to the 25^th^ and 75^th^ percentiles, respectively. Red and blue dots in violin plots indicate each data point. ***p < 0.001 **(C)** Stable value learning task. Four objects were always associated with a reward (high-valued objects), while the other 4 objects were always associated with no reward (low-valued objects). **(D and E)** Free-viewing tasks for testing visual habits. In the free-viewing task with 2 objects, two objects were selected pseudo-randomly from one set and presented simultaneously (D). In the free-viewing task with 4 objects, 4 objects appeared simultaneously (E). Monkeys were free to look at these objects (or look elsewhere) for 2 s without any reward feedback. ***p < 0.001

Next, for studying stable-value processing in the striatum, we first tested whether the monkeys exhibited habitual saccades based on previously learned stable values (Figures 1C and D) (*2*, *21*, *35*). The monkeys learned the values of fractal objects stably associated with reward during the object value-learning task (Figure 1C). After a retention period of >1 day following the last learning sessions, we examined their habitual saccades using the free-viewing tasks with 2 or 4 learned fractal objects (Figures 1D and E). In the free-viewing tasks, monkeys could freely look at the objects, although no rewards were delivered. After over 4 days of object-value associative learning, the monkeys shifted their gazes toward previously learned high-valued objects rather than low-valued ones (P < 0.001, two-tailed paired t-test) (Figures 1 D and E). Our data demonstrated that monkeys made habitual saccades guided by the stable object values they had previously learned, irrespective of the current reward outcome.

### Distinct value encoding in two example neurons of the putamen and ventral striatum

We recorded single-neuron activities across three striatal regions as the monkeys performed both the object value-reversal task and passive viewing task to examine how these individual neurons encoded flexible and stable values, respectively (Figures 2A and B). Figure 2 illustrates how the activity of two example neurons recorded from putamen (PUT) and ventral striatum (VS) was modulated in response to flexible and stable values. In the object value-reversal task, the activities of both PUT and VS neurons were modulated by changes in object-reward contingencies. Notably, these neurons exhibited a stronger response to high-valued objects when compared to low-valued objects (P < 0.001, Wilcoxon rank-sum test) (Figures 2C and E).

**Fig. 2.**
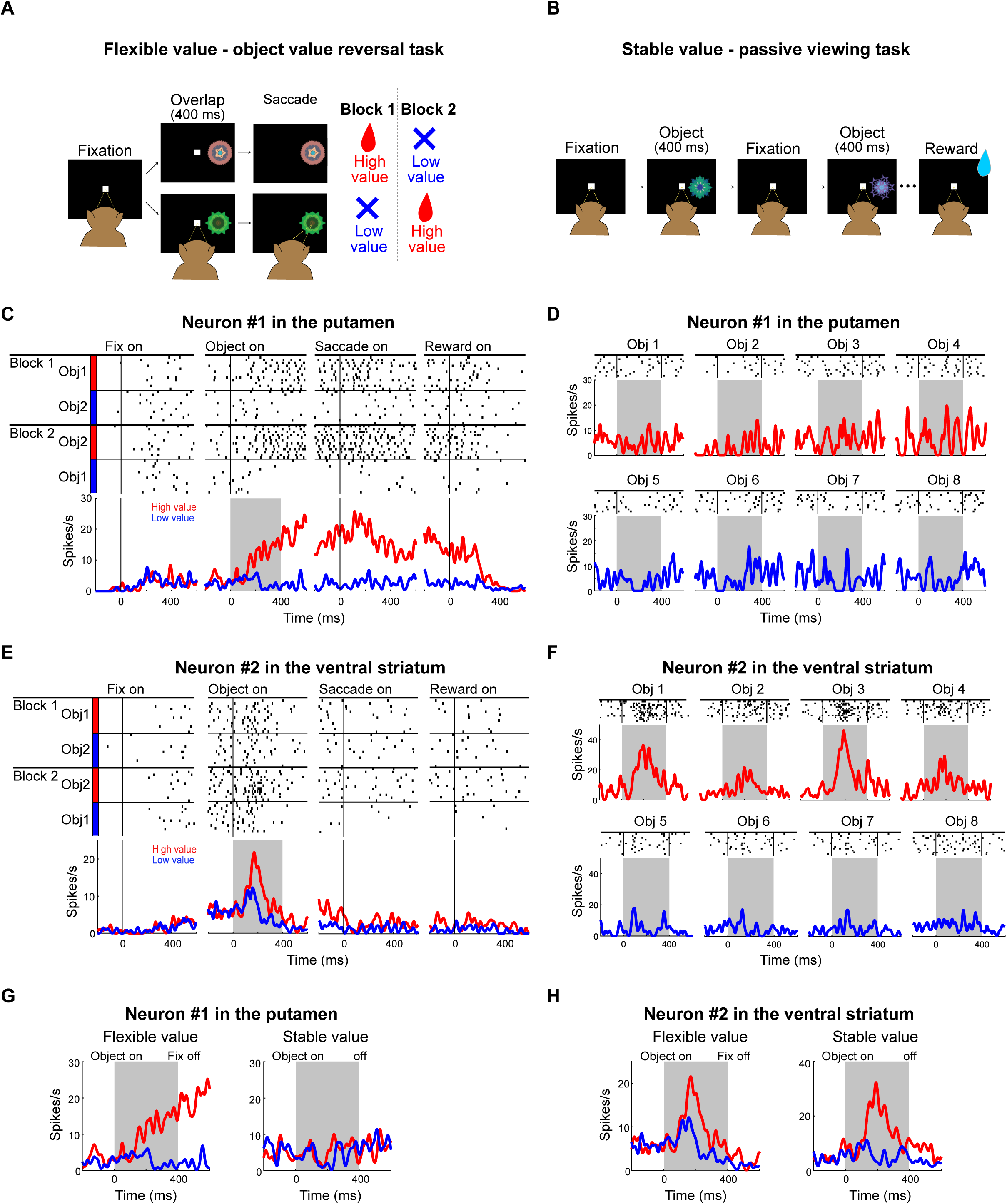
Example activities of PUT and VS neurons encoding flexible and stable values. **(A)** Object value-reversal task to examine the neural responses to objects with flexible value (same as in Figure 1A). **(B)** Passive viewing task to examine the neural responses to objects with a stable value. While the monkeys were fixating on a central white dot, two to six fractal objects were selected pseudo-randomly from a set of 8 objects and presented sequentially. The reward was delivered while the monkeys fixated after the last object disappeared. **(C and D)** A representative PUT neuron that exclusively encoded flexible value. **(C)** The responses of the representative PUT neuron to fractal objects are associated with flexible values. **(D)** The responses of the same PUT neuron to fractal objects are associated with stable values. **(E and F)** A representative VS neuron encodes both flexible and stable values. **(E)** The responses of the representative VS neuron to fractal objects are associated with flexible value. **(F)** The responses of the same VS neuron to fractal objects are associated with a stable value. **(G and H)** Averaged responses of the example VS and PUT neurons to fractal objects in object value-reversal task and passive-viewing task.

The responses of the same neurons to the stable value were examined using the passive viewing task. In this task, previously learned fractal objects were sequentially presented without any reward outcomes while the monkeys fixated on a central white dot (Figure 2B) (*2*, *17*, *21*, *35*). While both the example neurons recorded from PUT and VS exhibited activities encoding the flexible value, and their responses to the stable value were different. The PUT neuron did not exhibit activities encoding the stable value, whereas the VS neuron displayed activities encoding the stable value by responding more strongly to previously learned high-valued objects relative to the low-valued objects (Figures 2D and F). This example demonstrates that the PUT neuron selectively encoded the flexible value, whereas the VS neuron encoded both flexible and stable values (Figures 2G and H).

### Flexible value predominantly encoded in the putamen neurons

We identified individual neurons responding to visual objects during either the flexible or stable value testing procedures and then classified these neurons as object-responsive neurons. Among the object-responsive neurons identified in each structure, 251 out of 364 (68.96%) PUT neurons, 108 out of 137 (78.83%) caudate (CD) neurons, and 143 out of 187 (76.47%) VS neurons exhibited value discrimination responses in either or both flexible and stable value procedures (referred to as value-coding neurons). Neurons, whether encoding value or not, showed similar spike shapes across the three structures, suggesting that the recorded neurons were likely medium spiny neurons—a predominant type in the striatum (Figure S2A). The baseline activity of value-coding neurons was significantly lower than the non-value-coding neurons in the PUT and VS, albeit this difference in the baseline activity was not observed in CD (P < 0.001 in PUT, P < 0.05 in VS, Wilcoxon rank-sum test) (Figure S2B).

A total of 232 neurons (92.43%) out of 251 value-coding neurons encoded the flexible value and these neurons were widely distributed in all areas of PUT (Figure 3A). Among these neurons that responded to the flexible value, 133 neurons (57.33%) displayed high responses to high values (positive value-coding), while 99 neurons (42.67%) demonstrated high responses to low values (negative value-coding) (Figures 3B, S3A, and S3D). They distinguished flexible values by firing higher for the preferred values than for the non-preferred values (Figure 3C). Notably, most neurons (212 out of 251, 84.46%) exclusively encoded the flexible value (flexible value-coding neurons), as seen for an example neuron in Figures 2C and D. Neurons that exclusively encoded the stable value (stable value-coding neurons) or responded to both stable and flexible values (both value-coding neurons) were rare in PUT (Figures 3A and B). Only 7.57% of neurons (19/251) selectively encoded the stable value and 7.97% of neurons (20/251) encoded both values (Figure S3G).

**Fig. 3.**
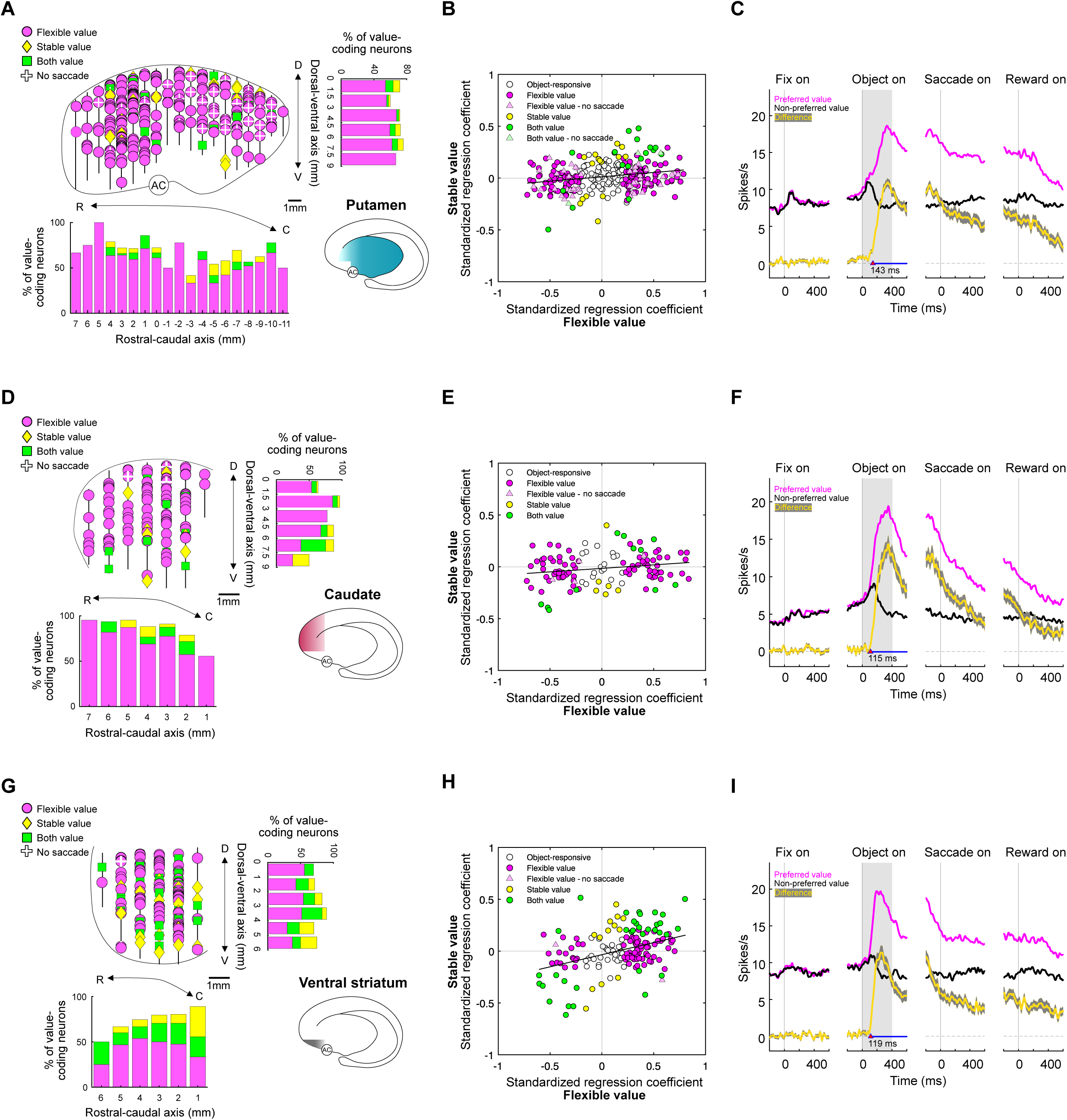
Population activity of flexible and stable value coding in the three striatal structures. **(A)** The locations of value-coding neurons in PUT in the sagittal view. D, dorsal; V, ventral; R, rostral; C, caudal. Neuronal distributions along the rostral—caudal axis and dorsal—ventral axis. In the rostral-caudal axis, 0 mm indicates the position of the anterior commissure (AC). In the dorsal–ventral axis, 0 mm indicates the dorsal border of PUT. The black vertical lines show the locations penetrated by electrodes. **(B)** Scatterplot of standardized regression coefficients for the stable value (ordinate) against those for the flexible value (abscissa) in PUT (n = 295). **(C)** The population response of PUT neurons that encoded flexible values during the object value-reversal task (n = 232). Neural responses are aligned to the fixation onset, object onset, saccade onset, and reward onset. Neuronal responses were averaged for the neurons’ preferred values (magenta) and non-preferred values (black). The yellow line indicates the difference between the preferred and non-preferred responses (mean ±SE). The blue line in bold indicates times that show statistically significant value discrimination activity (Wilcoxon rank-sum test). The red arrowheads indicate the initiation time of value discrimination. **(D–F)** Locations of value-coding neurons in CD (n = 115), value scatterplot, and averaged response of flexible value-coding neurons (n = 101). The same format as described in panels A–C. **(G–I)** The locations of value-coding neurons in VS (n = 164), value scatterplot, and averaged response of flexible value-coding neurons (n = 125). The same format as described in panels A–C.

### Putamen neural activity primarily modulated by value rather than saccades

Considering previous studies that demonstrated the role of PUT neurons in controlling motor movement, we investigated whether the activities of PUT neurons primarily contribute to object value processing or saccadic eye movements (*36–40*). To eliminate any potential influence of movement-related responses, we employed the no-saccade version of the object value-reversal task (Figure S4A). This task retained the same structure as the standard object value-reversal task but required the monkeys to maintain fixation at the center point without making saccades. Among the 183 object-responsive neurons recorded during the object value-reversal task without saccades, 92 neurons (50.27%) were flexible value-coding (Figure S4A). This observation was further supported by the results of our multiple linear regression analysis containing 341 object-responsive neurons recorded in the standard object value-reversal task (Figure 2A). Out of the analyzed neurons, the discharge rates of 208 neurons (61%) were significantly correlated with the object values (Figure S4B). However, only 41 neurons (12.02%) displayed a statistically significant correlation with the saccade reaction times (Figure S4B). Our data suggest that the neural responses modulated during the object value-reversal task were primarily associated with the processing of object values rather than saccadic eye movements.

### Predominant encoding of flexible value in the rostral caudate nucleus

The neurons encoding the flexible value were predominantly found in the rostral CD, which is consistent with previous findings (Figure 3D) (*2*). Among 108 value-coding neurons in the rostral CD, 93.52% (101 out of 108) of neurons encoded flexible values (Figures 3D and E). These flexible value neurons encoded values in either a positive or negative direction: 47.52% (48 out of 101) neurons were positive value-coding, while 52.48% (53 out of 101) neurons were negative value-coding (Figure 3E). The averaged activities of these neurons demonstrated a clear value discrimination response after the object presentation (Figures 3F, S3C, and S3E). However, we only found 6.48% (7 out of 108) neurons that exclusively encoded the stable value and 10.19% (11 out of 108) neurons encoding both values (Figures 3E and S3H).

### A relatively higher proportion of stable value-coding neurons in the ventral striatum

We found that 87.41% of value-coding neurons encoded the flexible value in the ventral striatum. However, unlike the dorsal striatum regions, 74.4% (93 out of 125) of neurons exhibited stronger responses to high-valued objects when compared to low-valued ones, while 25.6% of neurons displayed the opposite pattern (Figure 3H). The population of flexible value-coding neurons fired more in response to the preferred values than to the non-preferred values after the object presentation (Figures 3I, S3C, and S3F).

Interestingly, we found that VS showed a relatively higher proportion of neurons responding to the stable value (37.06%, 53 out of 143) when compared to the other structures of the striatum (Figures 3G and S3G-I). Moreover, 18 out of 53 neurons exclusively encoded the stable value, while the other 35 neurons encoded both values (Figures 3I and S3I). The proportion of these neurons tended to be greater in the caudal part than in the rostral part of VS, as previously described (Figure 3G) (*35*).

One notable observation was that, due to the predominant encoding of flexible values in CD and PUT, the coefficient values were dispersed widely along the flexible value axis, but not along the stable value axis, resulting in fitted slopes of 0.07 for CD and 0.08 for PUT (linear regression, P < 0.05 for CD, P < 0.001 for PUT) (Figure 3B and E). In contrast, when examining the coefficient values of individual neurons in VS, the fitted line exhibited a steeper slope of 0.22 when compared to those for CD and PUT, reflecting the substantial presence of neurons encoding both stable and flexible values (linear regression, P < 0.001) (Figure 3H).

### Dynamic adaptation of value-coding neurons to value reversal in the three striatal structures

In the object value-reversal task, the highest demand for cognitive flexibility occurred immediately after the objects’ values were reversed. After the value reversal, the monkeys had to relearn the new associations between objects and rewards to adjust their behaviors accordingly. Thus, if the neurons played a role in guiding this behavioral adaptation, it was crucial for their activities to dynamically respond to the behavioral shifts following reversals.

To investigate how flexible value-coding neurons dynamically respond to value reversal, we analyzed the changes in the discharge rates of these neurons alongside the corresponding shifts in saccade reaction times after the value reversal (Figure 4). In all three striatal structures, positive value-coding neurons revealed a decrease in their firing rates as the current value was switched from high to low, which negatively corresponded with a shift in the saccade reaction times from fast to slow (Figures 4A-C). Conversely, the discharge rates of negative value-coding neurons in these striatal structures decreased in parallel with an increase in the saccade reaction time following the value reversal (Figures 4D-F). We also found that, as the saccade reaction time transitioned from slow to fast following a shift in the object value from low to high, the firing rates of positive value-coding neurons increased, whereas the rates of negative value-coding neurons decreased (Figure S5). These changes in the firing rates of flexible value-coding neurons were correlated with behavioral adaptations that occurred in response to that value reversal.

**Fig. 4.**
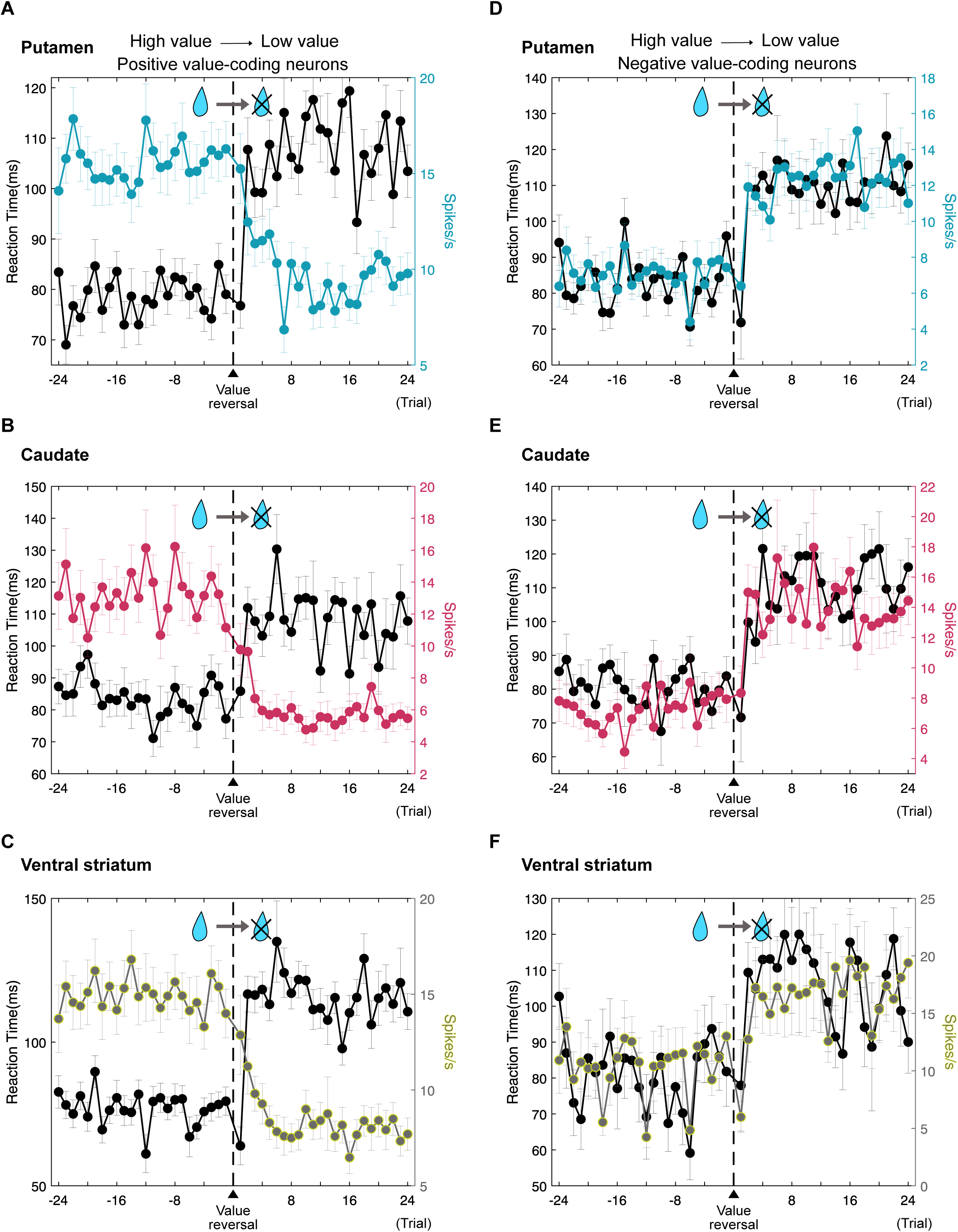
Dynamic changes in neural and behavioral responses to value reversal. **(A–C)** Changes in the response of positive value-coding neurons and saccade behavior before and after value reversal. Average firing rates (black dotted line) of positive-value coding neurons encoding flexible value in PUT (A), CD (B), and VS (C). The reaction times (red line for CD, cyan line for PUT, and yellow line for VS) are plotted as a function of the number of trials before and after the reversals from high value to low value. **(D–F)** Changes in the response of negative value-coding neurons and saccade behavior before and after value reversal. The same format as described in panels A–C.

Overall, our results demonstrated that a substantial population of neurons in the three structures of the striatum can dynamically adapt their responses in accordance with the changes in the object value. This observation suggests the crucial role of flexible value-coding neurons in PUT, CD, and VS for cognitive flexibility, a role that was not previously substantiated by the individual neural activities of PUT and VS during value reversal.

### Differential representation of task factors in three striatal structures

Next, we assessed the effects of the four task factors on object-responsive neurons in three striatal structures using a sliding-window regression analysis (Figure 5). The proportion of neurons representing object identity was significantly higher in CD and VS than in PUT between approximately 80 ms and 450 ms (Figure 5A). The neuronal proportions representing the object position in CD and PUT were significantly higher than the proportion in VS after object presentation (Figure 5B). The fraction of neurons representing reaction time was similar across the three structures (Figure 5C). In all structures, the effect of flexible value was the strongest among the effects of the four task factors (Figure 5D). The effect of value in CD was significantly stronger than that in VS and PUT across multiple time intervals. The value effects increased more slowly in PUT than in CD and VS. Thus, in addition to the object value, the three striatal structures represented other task factors, with variations in the intensity and latency of representation for each task factor among them.

**Fig. 5.**
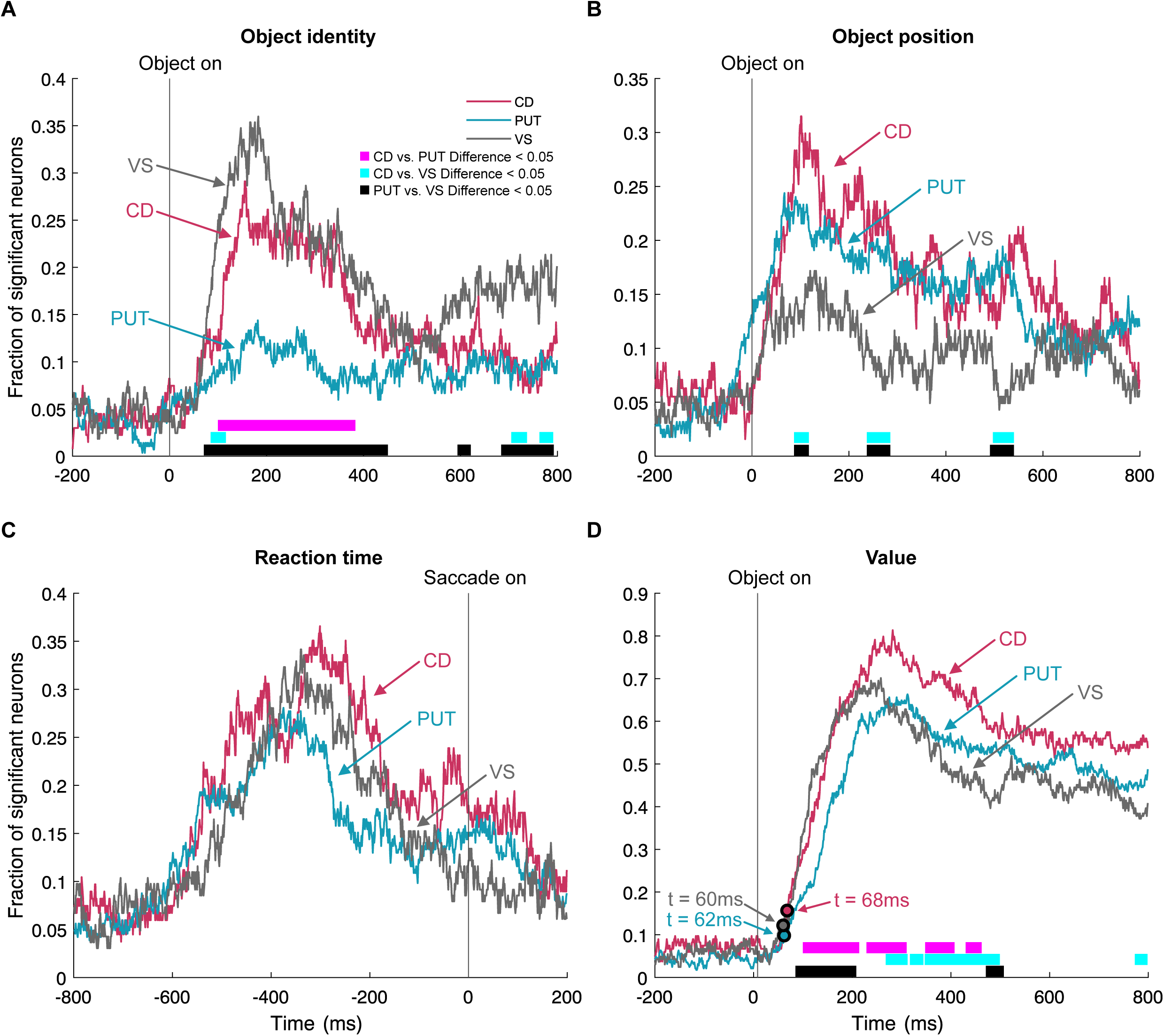
Representations of the task factors across the three striatal structures. **(A)** Time-varying fractions of object–responsive neurons are significantly modulated by object identity. The bars at the bottom indicate significant differences between all combinations of neural fractions from the three structures. **(B)** Time-varying fractions of object–responsive neurons significantly modulated by the object position. The same format as described in panel A. **(C)** Time-varying fractions of object-responsive neurons significantly modulated by the reaction time. The same format as described in panel A. **(D)** Time-varying fractions of object-responsive neurons significantly modulated by value. The same format as described in panel A. Arrows indicate times when the fraction of significant neurons exceeds the baseline levels.

### Faster value discrimination by the caudate nucleus compared to the ventral striatum

Previous studies have suggested serial processing of value information through these three structures (*41–43*). This serial value processing model is grounded in anatomical connections forming the ascending spiral between the three striatal structures and the substantia nigra in the primate basal ganglia (Figure 7B) (*44*). According to this model, the initial processing of value information occurs in VS neurons, followed by sequential processing involving CD and PUT neurons (*45–48*). To explore how these three structures differentially process value information according to the serial model, we compared the speed at which neurons in each structure encoded the flexible value. If this model works for the three striatal structures, we would expect to observe value encoding in VS first, followed by CD and PUT. However, we did not observe a meaningful faster encoding of the flexible value in VS neurons in comparison to that in CD and PUT neurons. In the averaged neural responses shown in Figure 3, the onset of value discrimination was nearly identical between CD and VS, and VS initiated value discrimination only 28 ms earlier than PUT (Figures 3C, F, and I). The onset times of value representation in the three striatal structures were very similar, as analyzed based on the fraction of neurons encoding value (ranges from 60 to 68 ms) (Figure 5D).

To further investigate the speed of flexible value encoding, we examined how many trials were necessary for neurons in each structure so as to exhibit value discrimination activity after value reversals. We incrementally added one trial at a time to determine the point at which the value discrimination activity reached statistical significance (Wilcoxon rank-sum test, p < 0.05). For example, a neuron in CD displayed statistically significant value discrimination activity within the initial 5 trials following the first exposure to the object values (Figure 6A). In contrast, an example neuron in PUT did not exhibit any significance until the 7th trial, and, by the 8th trial, the value discrimination activity reached statistical significance (Figure 6B). Similarly, an example neuron in VS displayed statistical significance in value discrimination activity in the 9th trial (Figure 6C). Notably, CD neurons exhibited flexible value encoding at the 6.68 ± 0.26th trial, which was significantly faster than the encoding in VS (Kruskal-Wallis test, Dunn-Sidàk test, p < 0.01) (Figure 6D). PUT and VS started encoding the flexible value at the 7.93 ± 0.28th, and the 8.61 ± 0.44th trials, respectively (Figure 6D).

**Fig. 6.**
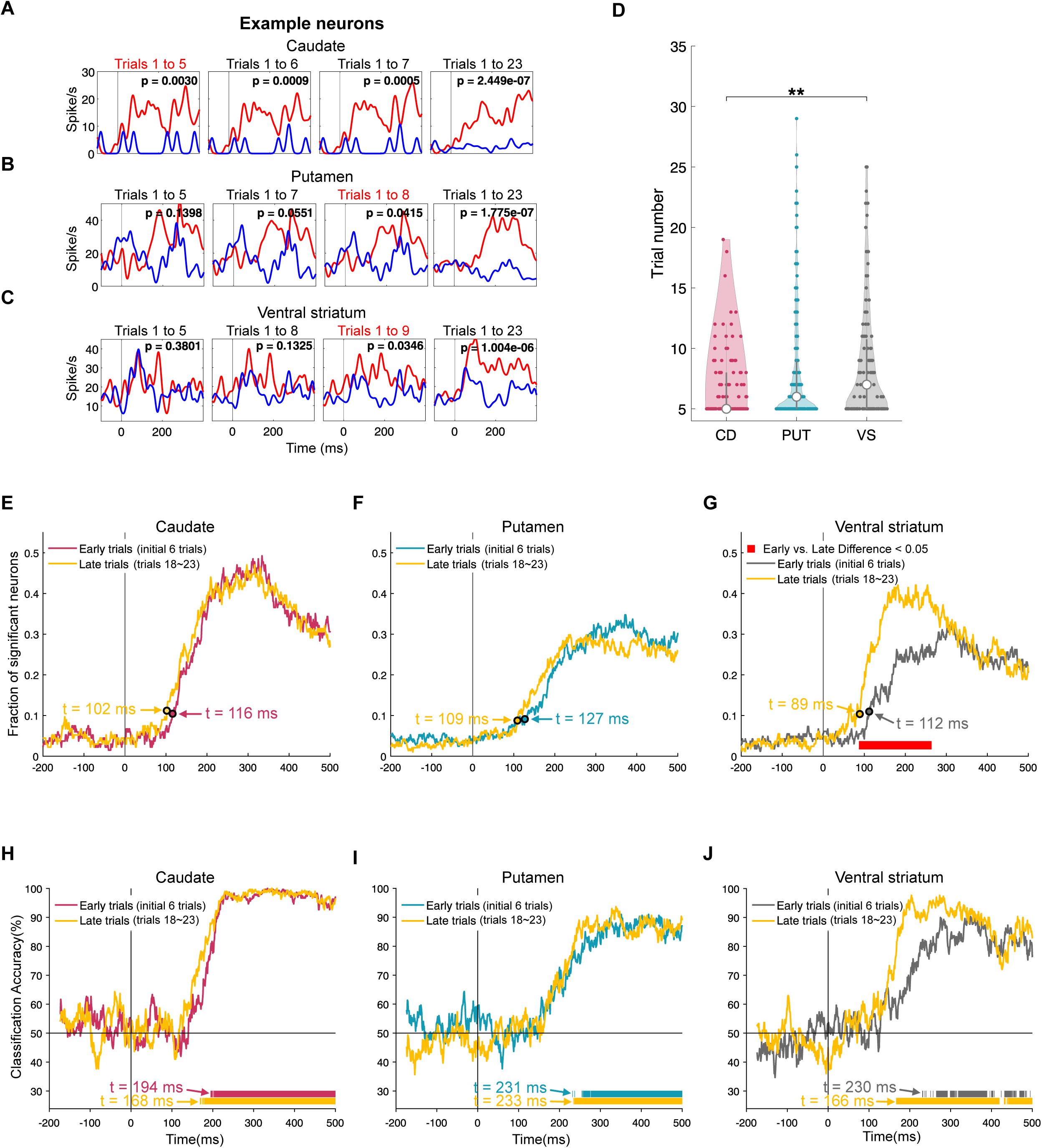
Speeds of value learning among the caudate, putamen, and ventral striatum. **(A– C)** Peristimulus histograms (PSTH) of example neurons in the three striatal structures that required different numbers of trials to discriminate values. The red “Trials 1 to number” highlights that the value discrimination activity of a neuron reaches statistical significance at that specified trial (p < 0.05, Wilcoxon rank-sum test). The black vertical line in a PSTH indicates the object onset, aligning with 0 ms. **(D)** Violin plots showing the number of trials required by neurons significantly encoding the flexible value in CD, PUT, and VS. The central white dot in a violin plot indicates the median, and the bottom and the top of the middle gray box correspond to the 25^th^ and 75^th^ percentiles, respectively. Small dots in violin plots indicate each data point. *p < 0.05, **p < 0.01 **(E–G)** Time-varying fractions of object–responsive neurons in CD (E), PUT (F), and VS (G) that significantly encoded the flexible value during the early and late trials. Arrows and “t” represent times at which the fraction of significant neurons exceeded the baseline levels. The red bar at the bottom indicates significant differences between the fractions of neurons in the early and late trials (G). **(H–J)** Comparison of decoding accuracies for flexible object values between the early and late trials. Arrows and “t” represent times at which the decoding accuracies were above the chance levels. Solid bars at the bottom of the plots depict the decoding accuracies for the late trials and early trials above the chance levels.

### Similar onset timings for initial value encoding across three structures

Several studies have demonstrated that VS neurons may play a critical role, particularly during the initial learning of object-value associations (*48–51*). We thus examined the value-learning responses of the three striatal structures during the early trials immediately after value reversals to analyze their differences in the onset times of value encoding (Figures 6E–J).

We compared the timings at which the fractions of neurons encoding the flexible value exhibited a significant increase over the baseline during the early learning trials (initial 6 trials after the value reversal). However, the onset times for encoding the flexible value were similar among these structures. The fractions of significant neurons in CD and PUT significantly increased at 116 ms and 112 ms, respectively, and the fraction in PUT displayed a slightly slower increase at 127 ms (Figures 6E–G).

To further identify the timing differences at the level of neural response pattern, we conducted a population decoding analysis using a maximum correlation coefficient classifier (Figures 6H–J). The neural population in CD exhibited the quickest onset of value representation at 194 ms during early learning. The onset times of value representation in VS and PUT were similar, occurring at 230 and 231 ms during the initial learning, respectively.

### Slow learners in the ventral striatum during reversed value acquisition

It is noteworthy that, during the early trials, the fractions of flexible value-coding neurons in PUT and VS were significantly lower than that in the fraction in CD (chi-squared test, p < 0.05, FDR corrected) (Figures 6E–G and S6A). However, when analyzed with all trials, there was no significant difference between CD and VS during the increase phase of the fraction, as described in Figure 5D. This finding suggests a possibility that VS neurons gradually developed flexible value-coding responses as the learning process progressed into the later trials.

Indeed, we observed that the fraction of flexible value-coding neurons in VS exhibited a statistically significant increase in the late trials when compared to that in the fraction in the early trials, indicating the presence of “slow learners” among VS neurons (chi-squared test, p < 0.05, FDR corrected) (Figures 6G and S6B). In contrast, the fractions of neurons encoding flexible value in CD and PUT did not significantly change between the early and late trials (chi-squared test, p < 0.05, FDR corrected) (Figures 6E and F). We also found that the onset timing of encoding flexible value in VS neurons during late trials became faster than during the early trials, with the onset timing changing from 112 ms to 89 ms (Figures 6G and S6B). However, the onset timing in CD and PUT displayed less change in comparison to VS (Figures 6 E and F).

To further examine these differences between the early and late trials in the neural response patterns, we conducted a population decoding analysis. The results showed that the value classification accuracy in VS increased with the advancing value learning (Figure 6J).

Notably, the onset timing of the classification accuracy in VS during late trials became faster than during early trials (230 ms during early trials and 166 ms during late trials) (Figure 6J). Neurons in other structures showed minimal change in the value classification accuracy and a smaller shift in the onset timing when compared to VS (Figures 6 H and I).

Our findings revealed a subset of VS neurons that were characterized as slow learners, requiring more time to learn flexible values when compared to neurons in other structures. However, once they had acquired the value, these VS neurons responded rapidly in processing the flexible value.

## Discussion

Our results thus demonstrated that neurons in the rostral regions of the three striatal structures predominantly encoded the flexible value, dynamically updating their neural activities in response to changes in the visual stimulus-outcome associations. In the three structures of the rostral striatum, a minority of neurons was observed to encode stable values for visual habit. These findings imply the involvement of the rostral striatum, including the caudate, putamen, and ventral striatum, in cognitive flexibility rather than habitual stability. This perspective challenges the previously proposed functional division of the striatum along the medial–lateral axis. Our findings introduce a principle of cognitive flexibility habitual stability process along the rostral—caudal axis of the primate basal ganglia, which complements previous studies showing stable value encoding in the caudal caudate, putamen, and ventral striatum (Figure 7A) (*2*, *35*, *52*).

**Fig. 7.**
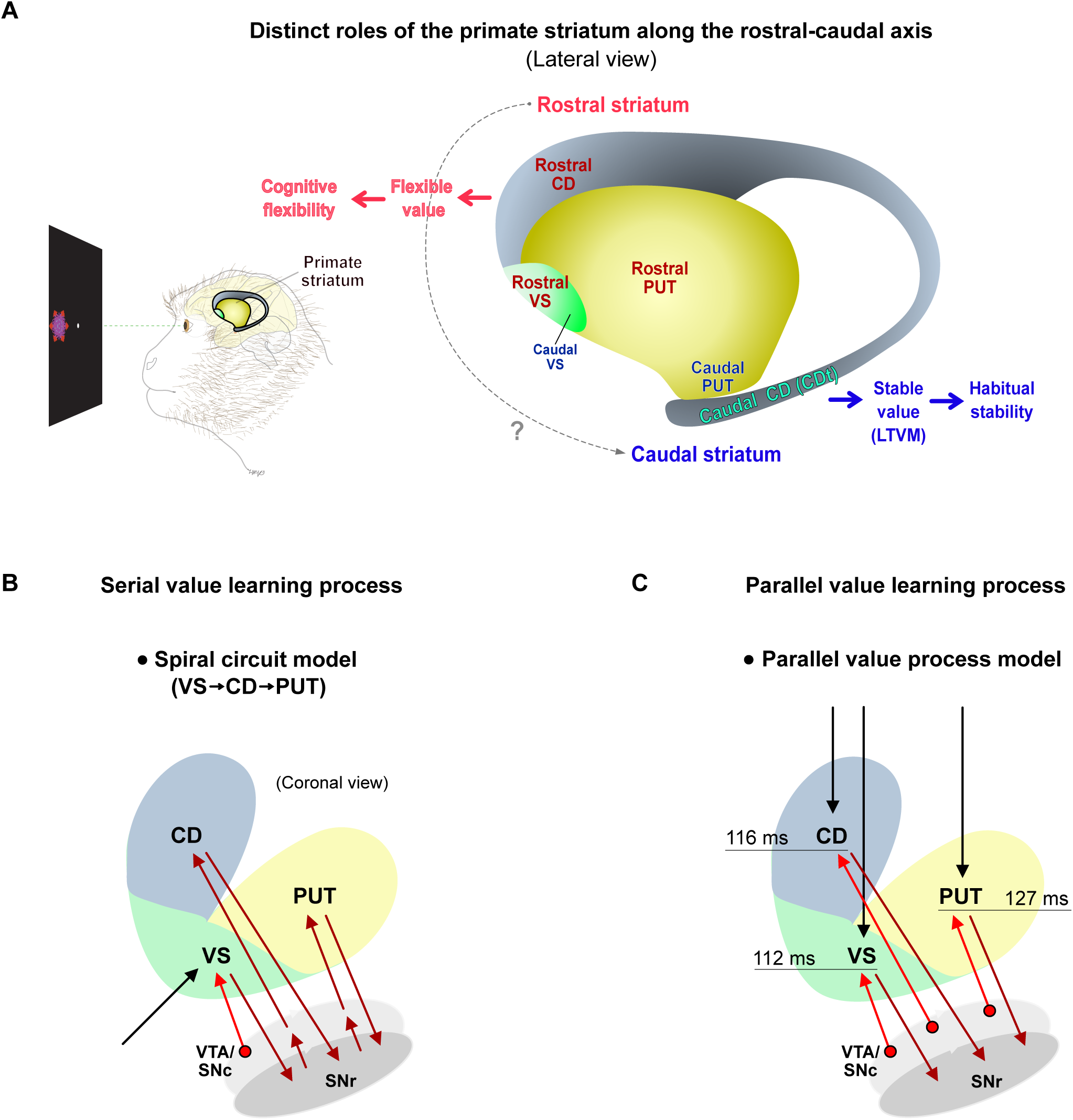
Models for processing of flexible and stable value in the primate striatum. **(A)** The functional division of the primate striatum along the rostral–caudal axis. The rostral striatum contributes to the encoding of the flexible value for cognitive flexibility, while the caudal striatum contributes to the processing of the stable value for habitual stability. **(B)** Spiral circuit model. In this model, VS serves as the initial learner of value information, transmitting this information to CD and then to PUT through the ascending spiral circuit between the striatum and dopamine areas (ventral tegmental area (VTA) and substantia nigra pars compacta (SNc)). **(C)** Parallel value process model. This model proposes that the three striatal structures learn object values independently by receiving RPE signals from dopamine neurons in VTA/SNc.

Furthermore, the observation that the rostral putamen processes cognitive flexibility suggests that it may not be the destination for value information guiding habits in an assumed serial value process (Figure 7B). Indeed, the speeds of value encoding were nearly identical across the three structures in the rostral striatum, implying parallel processing of flexible values in these rostral structures (Figure 7C). In addition, we identified a subset of ventral striatum neurons that learn values more slowly than neurons in the other structures, which challenges the previously proposed idea that the ventral striatum serves as the initial learner in cognitive flexibility processing.

### Cognitive flexibility and habitual stability encoding along the rostral–caudal axis of the primate striatum

Primate studies reported encoding of the stable value in the caudal regions of CD, PUT, and VS (*2*, *35*, *53*). The single-unit recording and inactivation studies showed that neurons in caudal CD stably maintained the value of objects and the inactivation of these neurons impaired visual habits (*2*, *16*). The caudal PUT, located adjacent to the caudal CD, shares the function of encoding stable values for habits with the caudal CD (*53*). In addition, previous human fMRI and single-unit recording studies showed that long-term value memory is more strongly encoded in caudal VS, as shown in our data (*35*).

Together with these previous findings, our results propose that the functional division of the primate stratum may align more with the rostral–caudal axis rather than the previously emphasized medial–lateral axis (Figure 7A). Notably, our findings highlight the engagement of the rostral regions of the three striatal structures in cognitive flexibility, while the caudal regions of these structures are implicated in habitual behaviors. Recent studies with humans and monkeys provide additional support for the functional distinctions of the primate striatum along the rostral–caudal axis (*11*, *13*, *54*). PUT inactivation impaired the performance of marmosets during the reversal task (*13*). Several human fMRI studies reported increased activity in rostral CD and PUT when behaviors were sensitive to the outcomes (*10*, *11*, *55*). However, as behaviors became habitual through extensive training, an increased BOLD signal was observed in caudal PUT (*54*). In addition, a comparative study between rodent and primate connectivity demonstrated that the caudal PUT, particularly, exhibited significant similarity with the DLS in rodents (*56*). Therefore, the DMS and DLS in rodents could potentially be considered homologous to the rostral and caudal striatum in primates, considering the parallel engagement of outcome-based processes in the rostral primate striatum and habitual processes in the caudal primate striatum.

### Does the value undergo a sequential processing from VS, CD to PUT in reversal learning?

Our neural recording data from across three striatal structures in each monkey revealed nearly identical onset times for flexible value representation in these striatal structures (Figures 5D and S6). The largest onset time difference between VS and CD was observed during late trials, amounting to only 13 ms. This finding suggests that the previously proposed transmission of value information from VS to CD and then to PUT through the ascending spiral circuit may need reconsideration (Figure 7B) (*44*, *45*).

The transmission of value information through this spiraling circuit, connecting the striatum and substantia nigra, requires a minimum of 45 ms. Specifically, approximately 30 ms are needed for the activation of VS neurons to impact the activity of neurons in substantia nigra pars reticulata (SNr), about 5 ms for activated SNr neurons to stimulate DA neurons in SNc, and roughly 10 ms for DA neurons in SNc to influence the activity of neurons in CD (*21*, *57*). In all the examined cases, the disparities in the onset times of value representation between the dorsal striatum (DS) and VS fell considerably short of 45 ms. This observation suggests that VS could not serve as the source of value information for the dorsal striatum in cognitive flexibility processing. This assertion gains additional support from the previous research, demonstrating that action-outcome learning can progress even in the presence of lesions in VS (*58*).

### The alternative model for value process: Parallel value process model

The model proposing serial value learning from VS to PUT through an ascending spiral circuit effectively elucidated a gradual transition from goal-directed to habitual behavior, aligning with previously proposed functional divisions within the striatum (Figure 7B) (*24*, *42*, *59*). In this conceptual framework, VS is in charge of the initial learning of outcome values as it is well-positioned to receive information about the current context, stimuli, and their affective values from the limbic system (*34*, *60*). The outcome values learned by VS are serially transmitted to CD and then to PUT through the spiraling dopaminergic circuit, driving the gradual transition from goal-directed behavior, mediated by CD, to habitual behavior, as governed by PUT.

The engagement of rostral regions across all three structures in cognitive flexibility challenges the necessity of this gradual transmission of value information. We accordingly proposed an alternative model that provides better explanations for the process of flexible value learning within the three striatal structures (Figures 7C). The parallel value process model suggests that the three striatal structures learn the flexible value of an object independently (Figure 7C). Considering that all three striatal structures receive RPE signals that are important in the learning process of outcome values, value learning in one structure does not need to rely on that of the other structures. Instead, all three striatal structures can independently learn the value of an outcome through each RPE signal. This model aligns with our data, as it does not presuppose that the learning and representation of value in a specific structure should occur earlier than the other structures both within a trial and across multiple trials.

### Value learning process across rostral and caudal striatum

Although the rostral striatal structures may learn the object value in parallel, the potential for a serial value learning process along the rostral–caudal axis of the primate striatum cannot be ignored. Anatomically, the ascending spiral circuit extends beyond the rostral striatum and persists into the caudal striatum, implying the possibility of serial transmission of value within the striatum from the rostral to caudal regions (Figure 7A) (*44*). Nonetheless, we cannot dismiss the possibility of a parallel value process in both the rostral and caudal striatum due to the presence of active closed-loop circuits between each striatal region and dopamine neurons (*57*). The next research question focuses on the mechanism underlying the learning of value across the rostral and caudal striatum toward understanding the complete process of value encoding at the primate circuit level.

## Methods

### Animals

Two female rhesus monkeys (Macaca mulatta, 5.0 and 5.2 kg, called NK and EV) were used. Animal care and experimental procedures were performed as approved by the Seoul National University Institutional Animal Care and Use Committee. Under general anesthesia and surgical conditions, a plastic head holder and a recording chamber were implanted into the monkeys’ skulls. The position of each chamber was tilted laterally by 25° such that it corresponded to the striatum. We started the training and recording sessions after the monkeys fully recovered from the surgery.

### Behavioral tasks

The behavioral procedures were controlled by a custom-made Visual C^++^-based experimentation data acquisition system (Blip; available at http://www.cocila.net/blip/). The monkey sat in a primate chair, facing a frontoparallel monitor screen in a sound-attenuated and electrically shielded room. Visual fractal stimuli created using Fractal Geometry were approximately 8° x 8° in dimension. The mean luminance was equalized across the images using the Spectrum, Histogram, and Intensity Normalization and Equalization (SHINE) toolbox written with MATLAB. An infrared camera (Oculomatic Pro) was used to record the eye position with a sampling rate of 1 kHz.

#### 1. Procedure for testing cognitive flexibility

##### Object value-reversal task

To examine behavioral and neuronal encoding of flexibly changing object value, we performed a task in which the object-value contingency was reversed in each block of 24–30 trials (*2*, *61*). For each monkey, fixed sets of two fractal objects (36 sets for monkey EV and 15 sets for monkey NK) were used as the saccade target. In each trial, one of them was presented at a right or left position pseudo-randomly (15° from the center). The fixation dot was presented simultaneously with the fractal object and disappeared after a 400-ms overlap period for recording the neural responses to the objects. Following the disappearance of the fixation dot, the monkeys were required to select the presented object by making a saccade to proceed to the next trial. In a block of 24–30 trials, one of the objects was associated with a reward (high-valued object) and the other with no reward (low-valued object). In the next block, the object-reward contingency was reversed. At least four blocks were included in one experiment. The object value-reversal task for the monkey NK included choice trials wherein the two objects were simultaneously presented at the right and left positions, and the monkey had to choose between the two objects. If the monkey correctly selected the high-valued objects, a liquid reward was provided. The choice trial appeared in every fifth trial.

To investigate whether the neural responses were associated with the monkeys’ movements rather than the flexible value, we introduced a version of the object value-reversal task without any saccades. In this no-saccade version of the reversal task, the objects were presented in the center instead of the right or left position, which eliminated the need for the monkeys to make a saccade. Following the monkeys’ fixation on an object for 400 or 600 ms, the reward was delivered according to the current object-reward contingency.

#### 2. Procedure for testing habitual stability

##### Object value-learning task

To generate stable long-term value memory, monkeys viewed visual objects repeatedly in association with consistent reward outcomes and thereby learned their stable values (*35*). In each of these sessions and the following tasks, a set of 8 computer-generated fractals was employed as visual objects. While the monkey was fixating on a central white dot, one of the objects was presented at the ipsilateral or contralateral position (15° from the center). The central fixation spot turned off 400-ms later, and the monkey was required to make a saccade to the object. Half of the objects were associated with a liquid reward (high-valued objects), whereas the other half were associated with no reward (low-valued objects). A tone was presented with either outcome. One training session consisted of 96 trials (12 trials for each object). Each set of objects was learned in one learning session in a day. The same sets of objects were repeatedly learned with the same object-value associations across days (>4 days), while new sets of fractals were introduced for learning across days. The monkeys EV and NK learned 51 (416 objects) and 73 (584 objects) sets of objects, respectively. Different sets of fractal objects were employed in procedures for testing cognitive flexibility and habitual stability.

##### Free-viewing task

To confirm whether the monkeys displayed a visual habit based on stable object value, we tested free gazes during which no instruction was provided but the learned fractal objects were presented, as previously used for monkeys and humans (*2*, *35*). In a free-viewing task with four objects, four objects were selected pseudo-randomly and presented simultaneously in four symmetric positions (15° from the center) after fixating on a central white dot for 300 ms. In a free-viewing task with two objects, two objects were pseudo-randomly selected and presented simultaneously at the ipsilateral and contralateral positions (15° from the center) after fixating on a white dot for 300 ms. The monkey was free to gaze at them for 2 s without any reward outcome. Separated from the free-viewing trials, a reward-associated white dot was presented during half of the trials to maintain the monkey’s motivation. In the free-viewing task with four objects, the reward-associated white dot was presented at one of the eight positions, and, in the free-viewing task with two objects, this dot was presented at one of three positions (i.e., the top, middle, and bottom). The reward was delivered when the monkey held its gaze on the white dot for 600 ms.

##### Passive-viewing task

This task was used to examine how neurons responded to the previously learned objects, as previously described (*2*, *35*, *61*). While the monkey was fixating on a central white dot, two to six objects, selected pseudo-randomly from a set of eight learned objects, were sequentially presented for 400 ms each at 15° from the center. Each learned object was presented at least 10 times in one session. To maintain the monkeys’ motivation, the reward was delivered 400 ms after the last object disappeared. Thus, the reward was not directly associated with any object. The value-coding activity was recorded after long-term learning (>4 days) with a sufficient retention period (>1 day after the last learning session).

##### Single-unit recording

As a monkey performed a task, the activity of single neurons in the three striatal structures was recorded by using a general method. The recording sites were determined by means of a 1-mm spacing grid system with the aid of MR images (3T, Siemens) obtained along the direction of the chamber. The single-unit recording was conducted using a glass-coated electrode (Alpha-Omega). The electrode was inserted into the brain through a stainless-steel guide tube and advanced by an oil-driven micromanipulator (MO-974A, Narishige). Neuronal signals from the electrode were amplified, filtered (250 Hz to 10 kHz), and digitized (30-kHz sampling rate and 16-bit A/D resolution) by using a Scout system (Ripple Neuro, UT). Neuronal spikes were isolated online using a custom voltage-time window-discrimination software (BLIP, Laboratory of Sensorimotor Research, National Eye Institute National Institutes of Health [LSR/NEI/NIH], available at www.cocila.net/blip), with the corresponding timings detected at 1 kHz. The waveforms of individual spikes were collected at 50 kH.

##### Analysis of single neural activity Object response

To examine the neuronal object responses, we counted the number of spikes within the test and control windows for each fractal object in the passive-viewing task and object value-reversal task. The control window was defined as 200–0 ms before the object onset, and the test window was defined as 0–400 ms after the object onset. In some of the sessions of the no-saccade version of the object value-reversal task, where objects were presented for 600 ms, the test window was 0–600 ms after the object onset. To test whether the neuron had object responses, we compared the numbers of spikes between the control and test windows in individual trials for each object. The Wilcoxon rank-sum test was employed to test for statistical significance.

##### Value-coding activity

We performed the linear regression analysis to determine whether the discharge rates of individual neurons recorded during the test window, as previously described, were significantly related to the value of an object. The discharge rates during the test window (*Y*) were fitted by the following linear regression model (Equation 1).

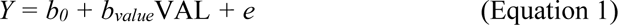

where VAL was the value of an object (1 and −1 for high value and low value); *b_value_* was the regression coefficient; and *b_0_* and *e* for the intercept and residual, respectively. If the *b_value_* was significantly different from zero (P < 0.05, t-test) and positive, we regarded the neuron as a positive value-coding neuron. If the *b_value_* was significantly different from zero (P < 0.05, t-test) and negative, we regarded the neuron as a negative value-coding neuron. Both positive and negative value-coding neurons were classified as value-coding neurons.

##### Neural fraction analysis

The time courses of the fraction of significant neurons (all at p < 0.05) for each task factor in the object value-reversal task were investigated through sliding window regression analyses using a 100-ms window with a 1-ms step. All neural fraction analysis was performed on data of object-responsive neurons recorded during the standard version of the object value-reversal task (n = 134 for CD, n = 285 for PUT, n = 164 for VS). When the time courses of neurons significantly affected by object identity, position, and value were analyzed, the activity of individual object-responsive neurons was aligned to the object onset and fitted by the following multiple linear regression model (Equation 2):

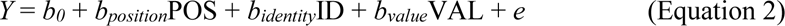

Where, POS is the object position (1 and −1 for right and left); ID is the object identity (1 and −1 for object1 and object2) VAL is the value of an object (1 and −1 for high value and low value); and b_position_, b_identity_, and b_value_ were the regression coefficient. We considered a neuron to encode a variable if the regression coefficient of that variable significantly differed from zero. After fitting this model to individual neurons, we had time courses of the fraction of significant neurons for each structure, considering each task factor. Significant differences in neural fractions between structures were assessed with a chi-square test. Initially, we performed a chi-square test at each time point to roughly investigate if there was a significant difference among the three structures. At p < 0.05 at a specific time point, we conducted additional pairwise chi-square tests at that particular time point to pinpoint where the significant differences existed—between CD and PUT, CD and VS, or PUT and VS. The raw p-values from this analysis suffer from multiple comparison problems as we applied the analysis across several time points and compared across the three structures. Therefore, we corrected for multiple comparisons using the false discovery rate (FDR) correction (*62*). For this purpose, the uncorrected raw p-values were sorted in ascending order. The rank-ordered p-values (P_(k)_) were considered significant when they were below the threshold defined by P_(k)_ ≤ (k/m)α for at least 6 consecutive bins. Where, k is the rank of the sorted p-values, α is the FDR, and m is the total number of tests (time points × _3_C_2_) under consideration. An α level of 0.05 was used for these tests.

When the time courses of neurons significantly influenced by the saccade reaction time were analyzed, the activity of individual object-responsive neurons was aligned to the saccade onset and fitted by the following multiple linear regression model (Equation 3):

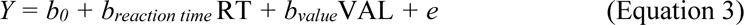

Where, RT is the reaction time and *b_reaction time_* and *b_value_* are the regression coefficients. The significant fraction differences among the three structures were analyzed by the same method mentioned earlier.

The fractions of neurons encoding value in the early and late trials were obtained by fitting the following model (Equation 4) to the activity of each object-responsive neuron, as given below:

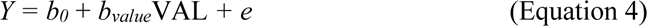

##### Onsets of value representation

To define the onset of value representation in neural fraction analysis, we compared the median of the baseline fraction (50–200 ms before the object onset) with the neural fraction encoding value at each time point (*62*, *63*). Statistical significance was tested by chi-square test, and the onset was defined as the beginning of 10 consecutive 1 ms steps that were significantly different from the median value (p < 0.05).

The onset of value discrimination in the averaged neural responses was investigated by a moving window analysis. The 1-ms window was moved by the 1-ms time bin, and the value discrimination activity (high-valued object – low-valued object activity) at each time point was compared to the baseline (0) by paired t-test (*64*). The onset was defined as the beginning of 10 consecutive 1-ms steps that were significantly different from the baseline value (p < 0.05).

##### Speed of value learning

The data of flexible value-coding neurons recorded during the standard and no-saccade versions of the object value-reversal task were employed to examine the value-learning speeds of neurons in the three striatal structures (n = 121 for CD, n = 278 for PUT, n = 128 for VS). To examine how many trials were required for flexible value-coding neurons in each structure to exhibit value discrimination activity after the initial exposure to object value, we first aligned the trials in each block (4 blocks in a session) from the first trial to the last trial. Then, we examined whether a neuron responded differently to high and low values during the test window by performing the one-tailed Wilcoxon rank-sum test on neural data from the initial 5 trials following the first exposure to object values within each block. The test window was defined as the timeframe during the fixation dot-object overlap period (0–400 or 0–600 ms after the object onset). In this analysis and the analysis comparing neural activities between the early and late trials (Figures 6E–J), the first trial in each block wherein a monkey first encountered the value of an object was designated as trial 0, with subsequent trials denoted as trial +1, trial +2, etc. By analyzing the data from the entire set of trials, we could discern whether a neuron fired more to a high or low value. This information was used to specify the direction of the one-tailed Wilcoxon rank-sum test. In addition, in most cases, the grouping of the initial 5 trials within each block (5 trials × 4 blocks = 20 trials) provided us with a dataset exceeding the minimum sample size required for conducting a one-tailed Wilcoxon rank-sum test (at least 3 samples for each group). However, in cases where the available data did not meet the minimum sample size requirement, we excluded the data of that specific neuron from this analysis. This exclusion constituted a minimal proportion, accounting for 0.0468% of the data recorded in CD, 0.0331% in PUT, and 0.0078% in VS. If a neuron did not exhibit significant value discrimination within these initial 5 trials, we incrementally added one trial at a time and performed the rank-sum test to determine the point at which the value discrimination activity reached statistical significance.

##### Analysis of neural population activity

###### Neural decoding analysis

To compare the efficiency of value representation between the three structures and between the early and late phases, we employed a maximum correlation coefficient classifier implemented in the Neural Decoding Toolbox (*65*). From 134 neurons in CD, 284 neurons in PUT, and 163 neurons in VS that responded to objects during the saccade version of the object value-reversal task, we randomly selected 90 neurons and created the pseudo-populations for each structure. The firing rates of individual neurons were sampled with a 30-ms bin and at 1-ms resolution. Trials were randomly split into training and test sets using 3-fold cross-validation, where the classifier was trained on data from 2 splits and tested on the 3^rd^ split, while the procedure was repeated thrice using a different test split each time. The classifier was trained on the firing rates of a neural population to discriminate the value of an object. The classification accuracy was then calculated as the percentage of predictions that were correctly made on trials from the test set. We repeated this procedure over 50 resample runs, where different random populations of training and test splits were created on each run. We averaged these resample runs and made one decoding accuracy.

To estimate when the decoding accuracy was above the expected by chance, we employed a permutation test. For this, we randomly shuffled the labels of each variable in each trial separately for each neuron. Next, we ran the same decoding procedure and made a null distribution of shuffled decoding. We repeated this procedure 7720 times and generated a full shuffled decoding null distribution for each time bin. The solid bars on the bottom of Figures 6H–J show the period when the decoding accuracy was >7720 points in the shuffled decoding null distribution (P < 1.2953e-04 for each time bin).

## Acknowledgments

This work was supported by the Neurological Disorder Research Program (NRF-2020M3E5D9079908) through the National Research Foundation (NRF) of Korea, and Korean government (MSIT) grant (NRF-2019R1A2C2005213).

## Author contributions

H.F.K. designed and supervised the entire project. SY. A., SH. H. and H.F.K. performed the behavior and single-unit recording experiments. SY. A. and SH. H. analyzed the data and prepared the figures. SY. A. and SH. H. wrote the first draft, and SY. A., SH. H., KW. L., and H.F.K. interpreted data and wrote the final manuscript.

## Data availability

All data needed to evaluate the conclusions in the paper are present in the paper and/or the Supplementary Materials.

The original/source data can be provided by Hyoung F. Kim pending scientific review and a completed material transfer agreement. Requests for the original/source data should be submitted to: Hyoung F. Kim.

## Supplementary figure legends

**Fig. S1.**
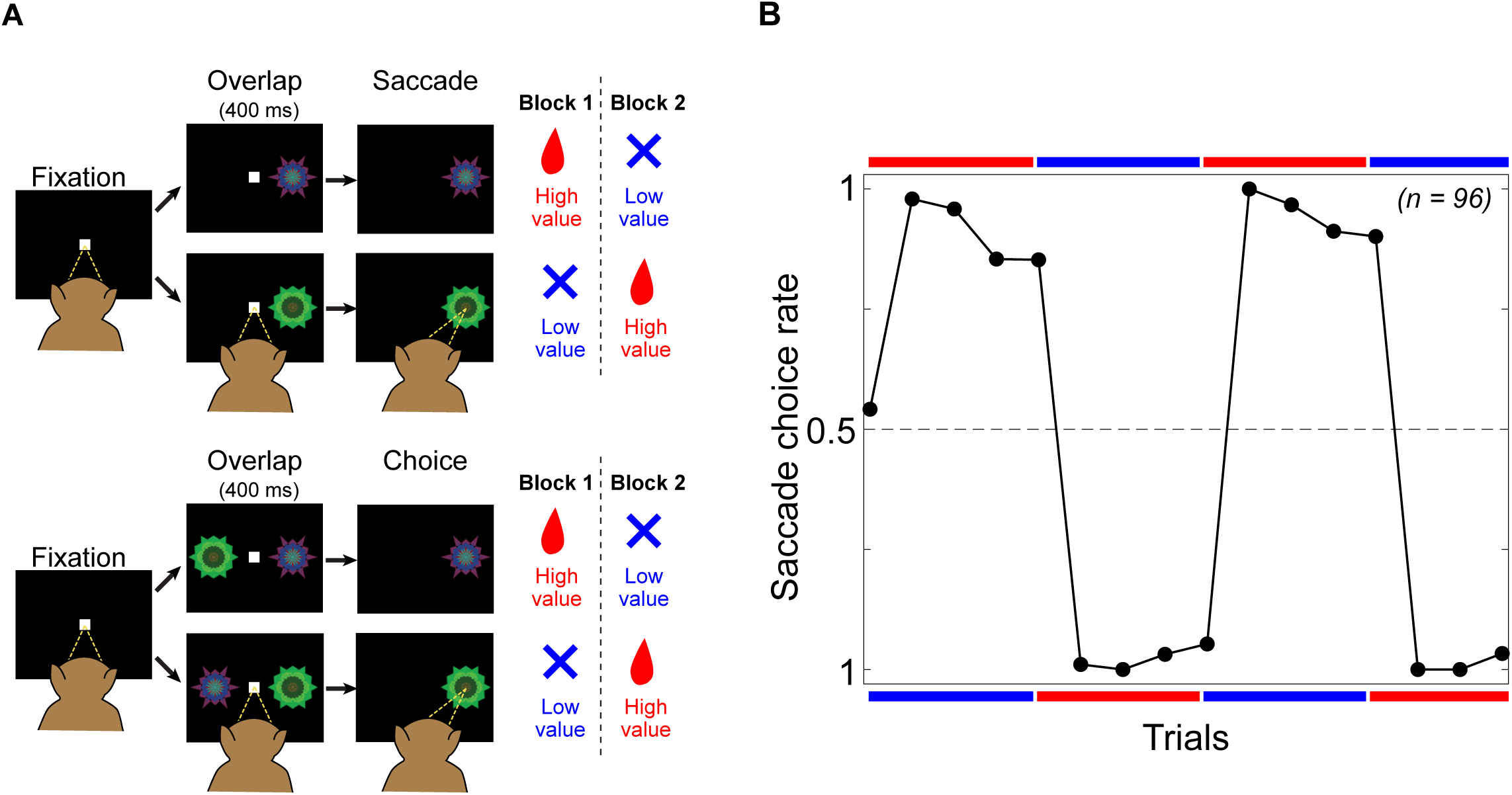
Saccade choice rates in choice trials of object value-learning task. **(A)** The sequence of events in the object value-reversal task including saccade trials (top, same as in Figure 1A) and choice trials (bottom panel). In the object value-reversal task with choice trials, a choice trial appeared at every fifth trial. High-valued and low-valued objects were simultaneously displayed on the screen and the monkey was required to choose one of them by making a saccade. **(B)** Flexible adaptation of the monkeys’ choice to the change in object values. The average choice rate for 96 sessions is plotted against each trial. The plots close to the horizontal red bars indicate choices preferring the high-valued object. The plots close to the horizontal blue bars indicate choices preferring the low-valued objects.

**Fig. S2.**
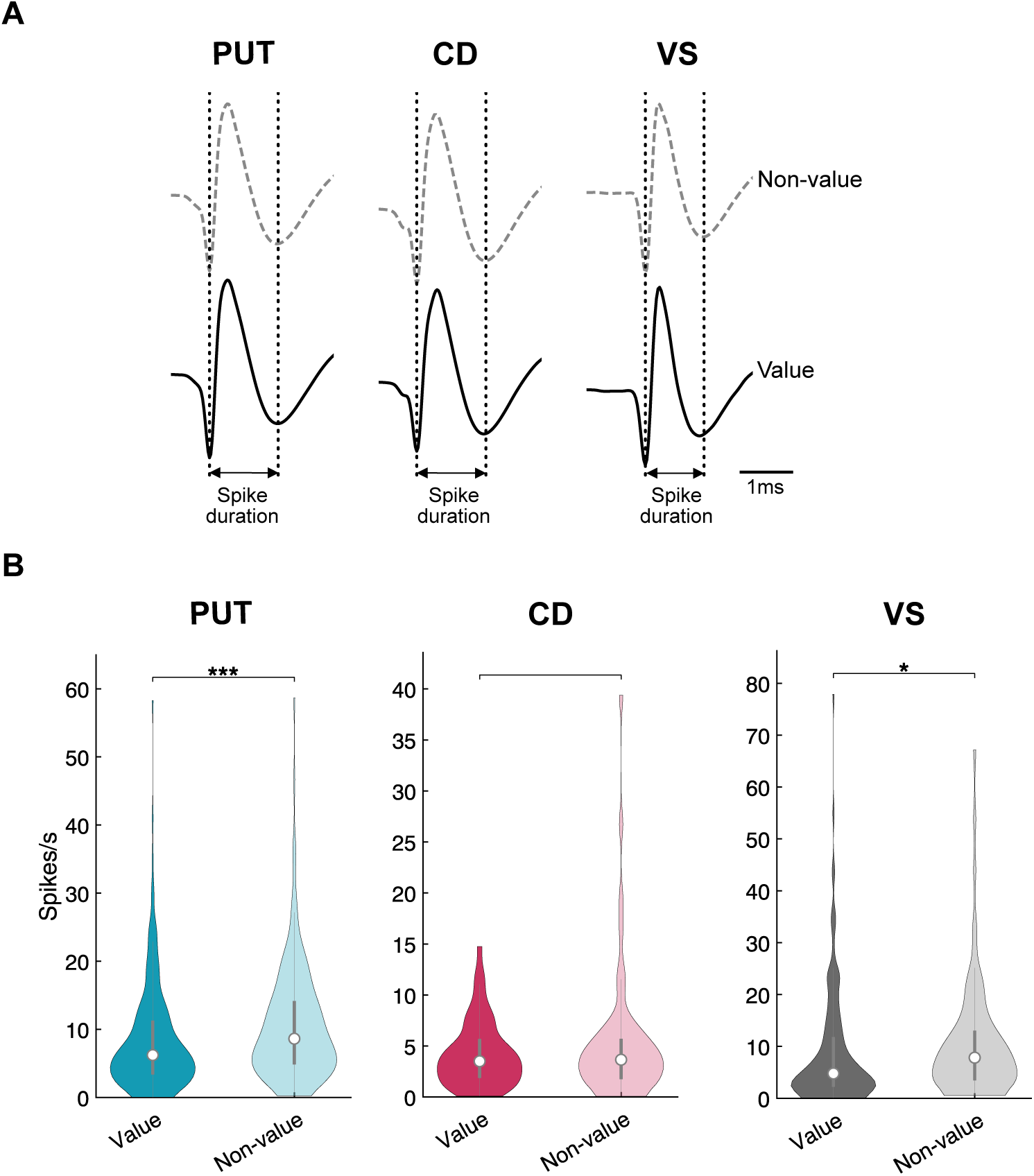
Electrophysiological properties of neurons in the three striatal structures. **(A)** Spike shapes of non-value-coding (dashed line) and value-coding neurons (solid line). In all three striatal structures, non-value-coding and value-coding neurons had similar spike shapes. **(B)** The baseline activities of non-value-coding and value-coding neurons. The baseline activity is the mean firing rate during 500 ms before the onset of the fixation dot (inter-trial-interval period). The central white dot in a violin plot indicates the median, while the bottom and the top panels of the middle gray box correspond to the 25^th^ and 75^th^ percentiles, respectively (**p < 0.01, ***p < 0.001, two-tailed t-test).

**Fig. S3.**
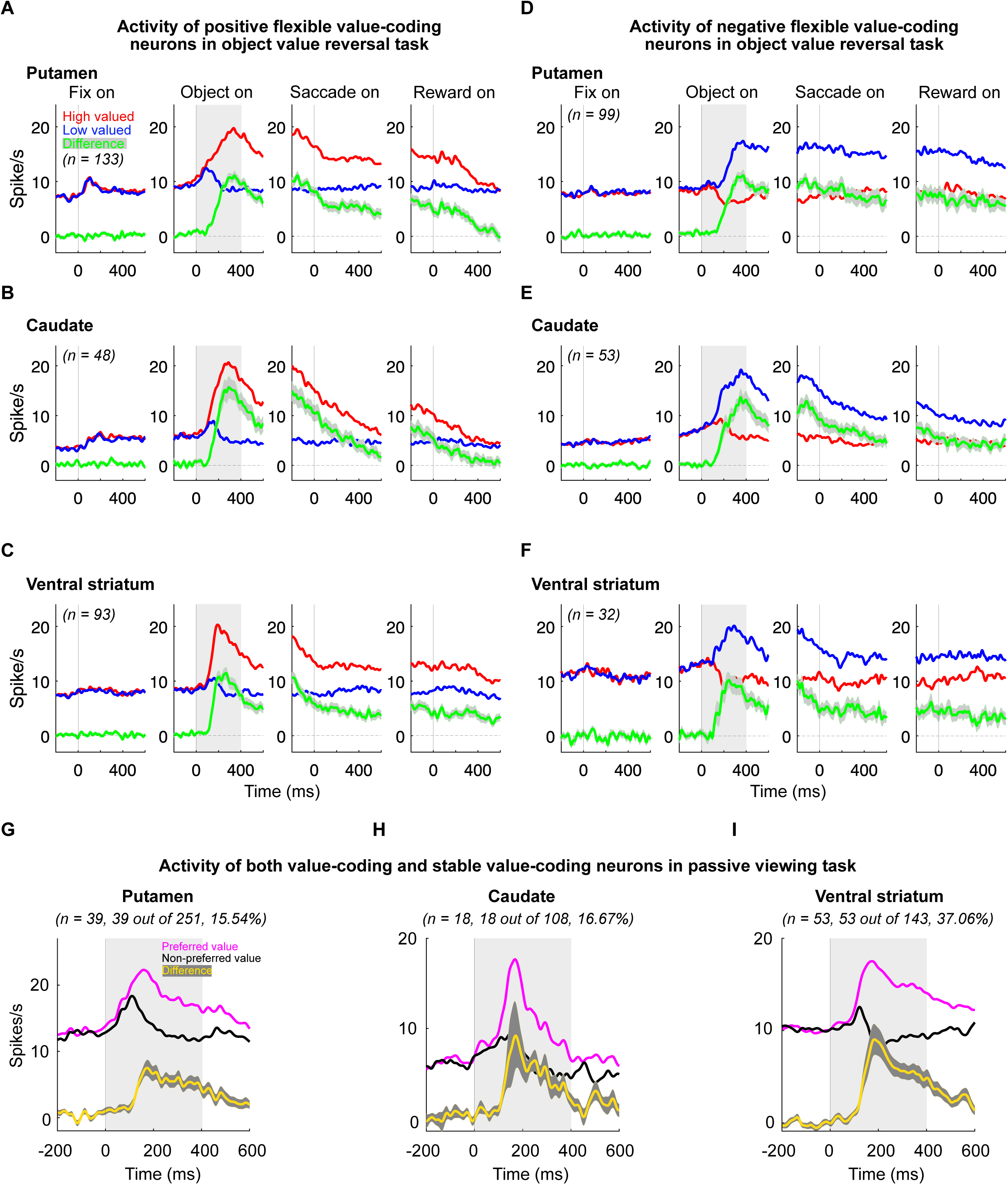
Positive and negative encoding of the flexible and stable values in neurons of the three striatal structures. **(A–C)** Population responses of neurons in PUT (A), CD (B), and VS (C) that positively encoded the flexible value. The number of neurons is indicated at the top left panel. **(D–F)** Population responses of neurons in PUT (D), CD (E), and VS (F) that negatively encoded the flexible value. **(G–H)** Averaged neural responses in PUT (G), CD (H), and VS (I) that encoded the stable value. **(G)** Among 251 value-coding neuros, 39 PUT neurons encoded the stable value. **(H)** Among 108 value-coding neuros, 18 CD neurons encoded the stable value. **(I)** Among 143 value-coding neuros, 53 VS neurons encoded the stable value.

**Fig. S4.**
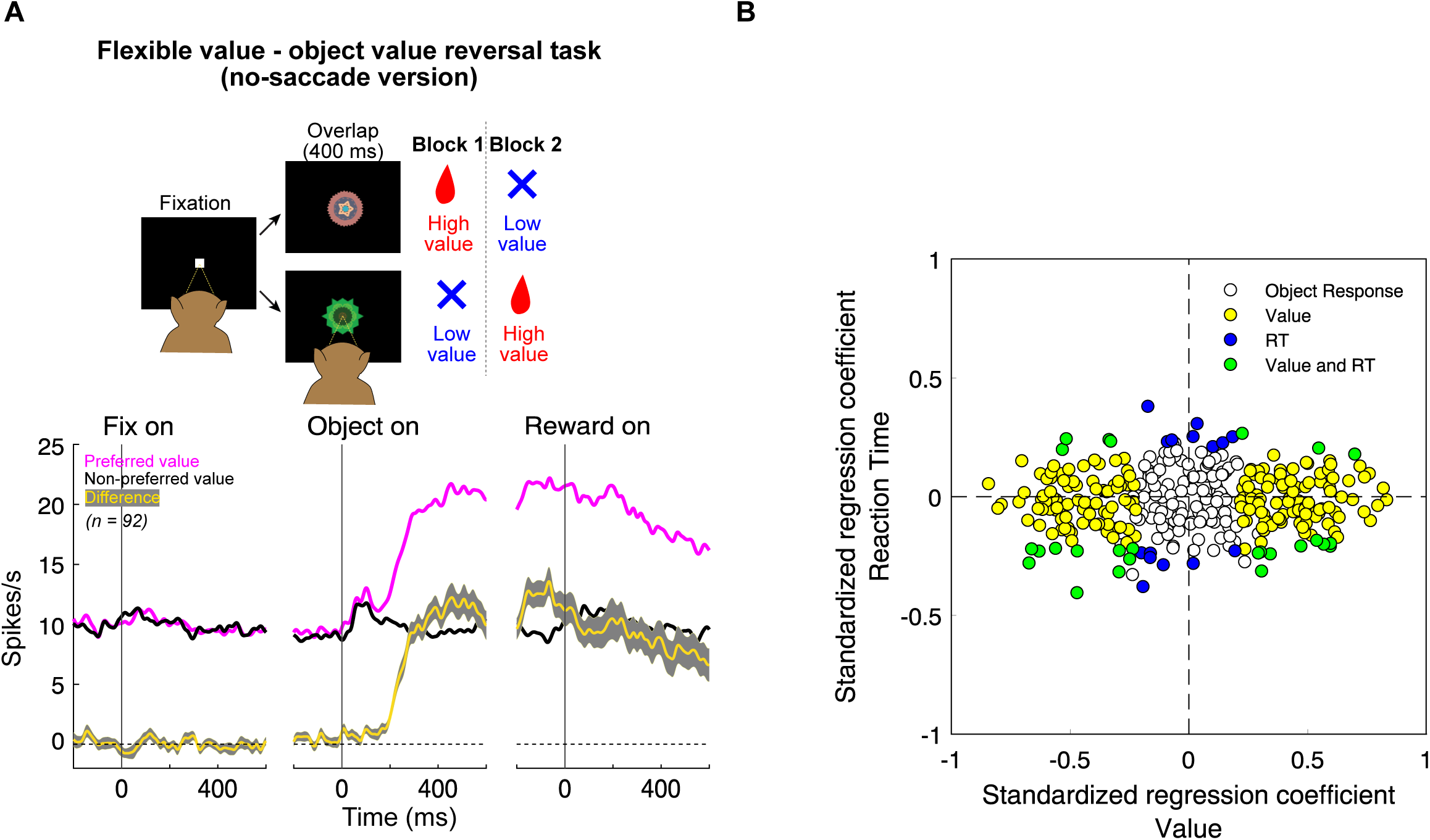
Predominant encoding of object value rather than saccadic eye movement in PUT neurons. **(A)** The no-saccade version of the object value-reversal task scheme and the population response encoding flexible value. The experimental scheme is depicted at the top panel of Figure A. While monkeys were fixating on a central white dot, one of two fractal objects was presented at the center. The monkeys were required to maintain fixation, with no saccade allowed. A liquid reward was delivered for one object, but not for the other after 400 or 600 ms of fixation. The object-reward contingency remained unchanged in a block of 24– 30 trials and then was reversed in the next block. The averaged PSTH of neurons that encoded flexible values in this task are shown at the bottom panel of Figure A (n = 92). **(B)** Scatterplot of standardized regression coefficients for the reaction time (ordinate) against those for the flexible value (abscissa) in PUT (n = 341).

**Fig. S5.**
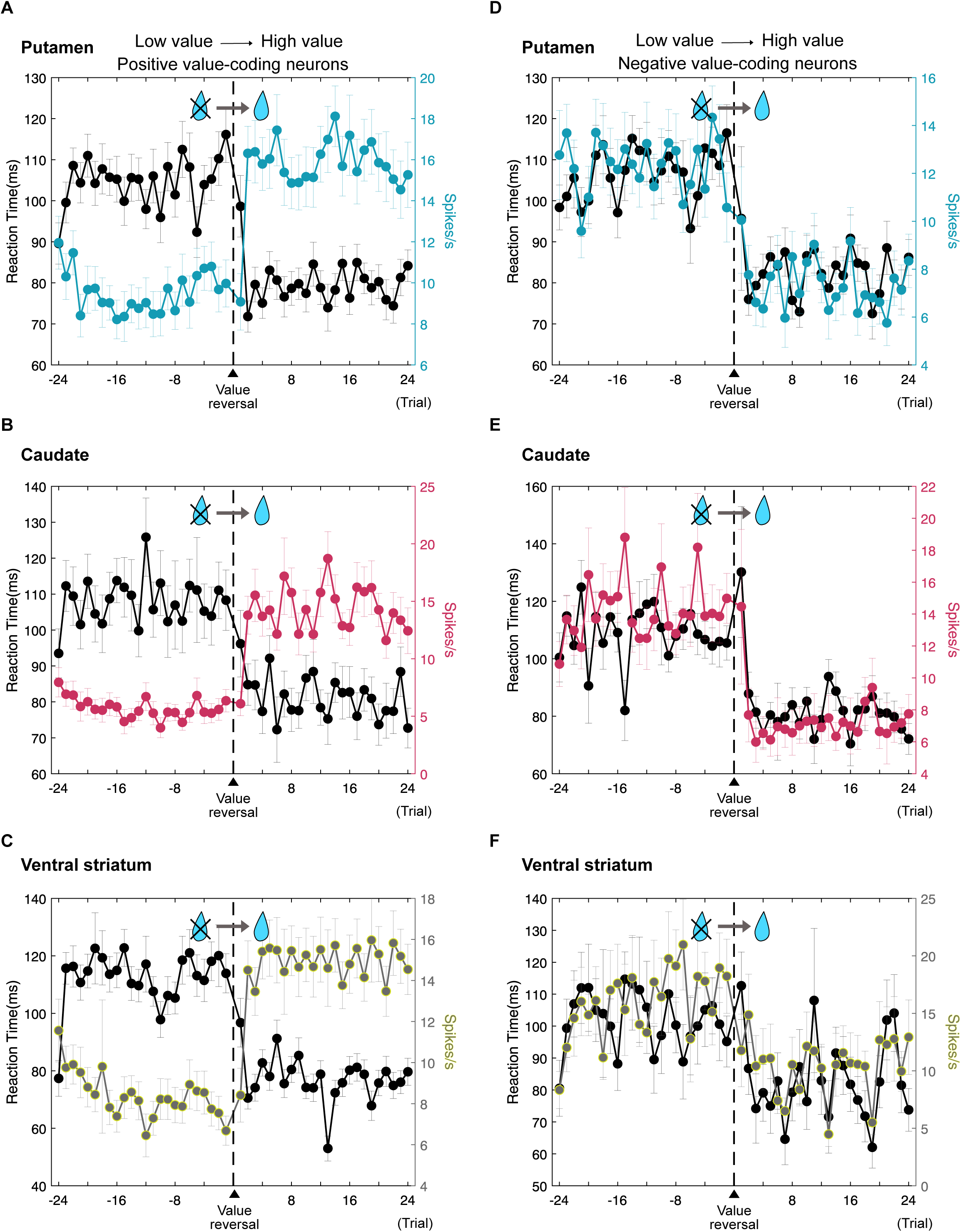
Dynamic alterations in neural and behavioral responses during the transition from low value to high value. **(A–C)** Changes in the response of positive value-coding neurons and saccade behavior before and after value reversal. The average firing rates (black dotted line) of positive-value coding neurons encoding flexible value in PUT (A), CD (B), and VS (C), and reaction times (red line for CD, cyan line for PUT, and yellow line for VS) are plotted as a function of the number of trials before and after the reversals from low value to high value. **(D–F)** Changes in the response of negative value-coding neurons and saccade behavior before and after value reversal. The same format described as in Sections A–C.

**Fig. S6.**
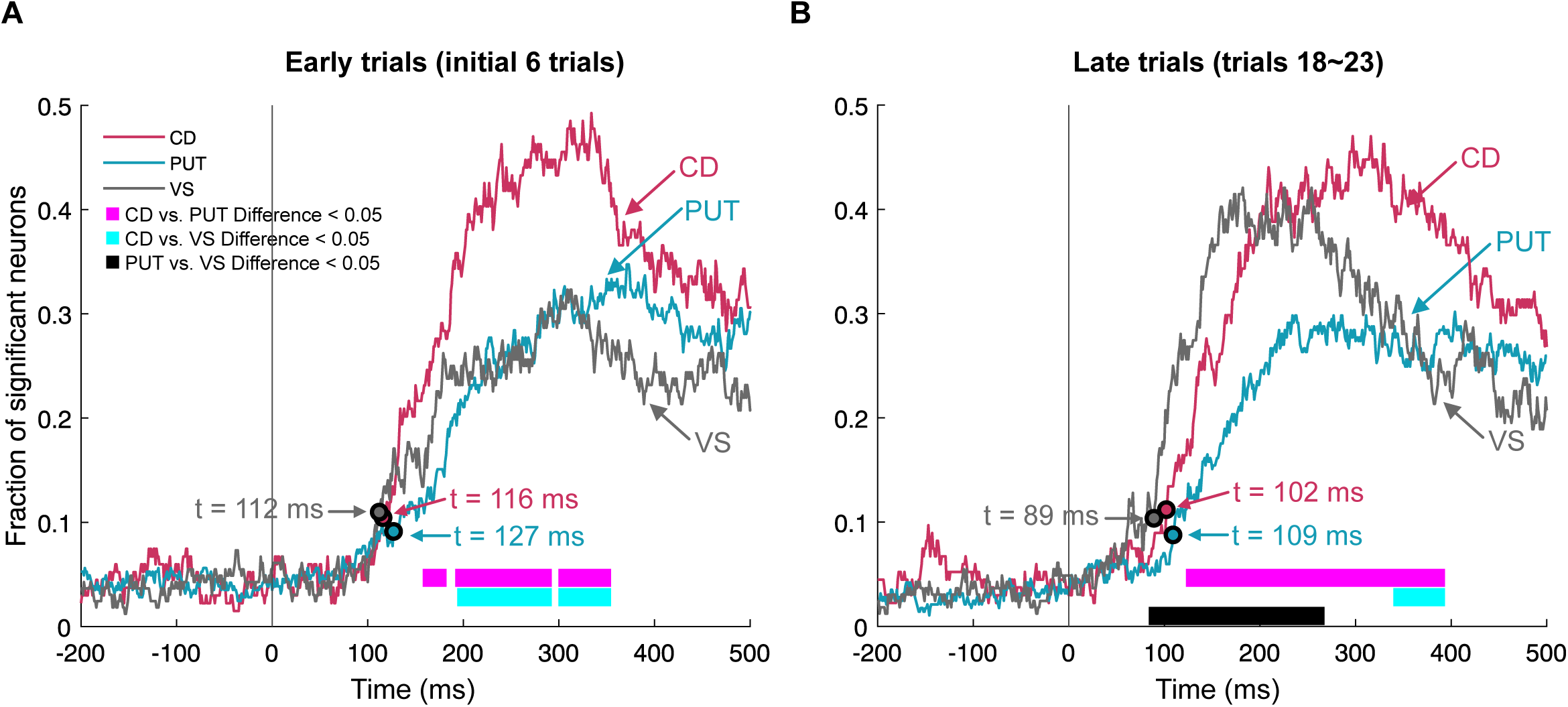
Representation of the flexible value in the three striatal structures during early and late stages of learning. **(A–B)** Time-varying fractions of object-responsive neurons in the three striatal structures significantly encoded the flexible value during early trials (A) and late trials (B). Arrows and “t” represent times when the fraction of significant neurons exceeds the baseline levels.

## Supplementary methods

We computed the saccade choice rate for high-valued objects using the following formula: nSACg/(nSACg + nSACb), where nSACg and nSACb are the numbers of saccades toward high-valued and low-valued objects, respectively.

When the influence of value and reaction time on the discharge rates of object-responsive neurons in PUT was examined, the discharge rates of object-responsive neurons in PUT during the 400-ms period after the object onset (*Y*) was fitted by the following multiple linear regression model (Equation 5):

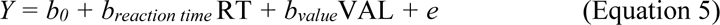

Where, RT is the reaction time; VAL was the value of an object (1 and −1 for high value and low value); *b_reaction time_* and *b_value_* are the regression coefficients; and *b_0_* and *e* for the intercept and residual, respectively. The discharge rates of a neuron and the reaction time were z-score standardized before being fitted by the aforementioned model.

